# Generalizable strategy for designing helical binders rigidly fused to large protein cages with disulfide bonds

**DOI:** 10.64898/2026.11.26.769080

**Authors:** Roman Aguirre, Hongbo Zhu, Todd Schurr, Robert Damoiseaux, Z. Hong Zhou

**Affiliations:** Department of Chemistry and Biochemistry, University of California, Los Angeles (UCLA), Los Angeles, CA 90095, United States; Department of Microbiology, Immunology, and Molecular Genetics, UCLA, Los Angeles, CA 90095, United States; The California Nanosystems Institute (CNSI), UCLA, Los Angeles, CA 90095, United States; Department of Molecular & Medical Pharmacology, UCLA, Los Angeles, CA 90095, United States

## Abstract

Self-assembling protein cages are versatile scaffolds for multivalent antigen presentation and single-particle cryogenic electron microscopy (cryoEM) of small proteins, yet adapting genetic fusions to diverse targets typically demands extensive construct-specific optimization. Here, we establish a modular display platform coupling *de novo*-designed α-helical binders to the trimeric subunit of two icosahedral protein nanoparticle cages: I53-50 and pyruvate dehydrogenase complex (PDC) E2p core. Using the 16 kDa first bromodomain of bromodomain-containing protein 4 (BRD4-BD1), a therapeutically relevant epigenetic target below the practical size limit for single-particle cryoEM, as proof-of-concept cargo, we generated and screened binder candidates *in silico* and *in vitro*, identifying a soluble binder with a steady-state dissociation constant (*K_d_*) of 13 ± 8 µM. Although continuous N-terminal α-helical fusion permitted high-yield assembly of 60-copy I53-50 and PDC E2p nanoparticles, cryoEM reconstructions revealed high core resolution (2.34 Å and 2.97 Å, respectively) alongside poorly resolved binder density caused by lever-arm amplification of small angular fluctuations at the fusion junction. PDC constructs were additionally limited by poor solubility, and subsequent work used I53-50 alone. To rigidify the interface, we engineered an inter-domain disulfide bond (C104–C213) across the junction and redesigned neighboring residues using ProteinMPNN. Molecular dynamics simulations indicated that disulfide stapling suppressed interfacial fluctuations, and cryoEM resolved the four-helix bundle binder at 3.2 Å and the BRD4-BD1-bound complex at 3.7 Å (3.0–4.0 Å local resolution). Combining target-specific helical binders with a reversible disulfide staple provides a plug-and-play architecture adaptable to cellular cryogenic electron tomography, redox-responsive drug delivery, and vaccine design.

## Introduction

Self-assembling protein nanoparticles exhibit diverse symmetric architectures formed through the precise oligomerization of one or more protein subunits. While many protein cages occur naturally as compartments for metabolic enzymes and biochemical reactions, engineered variants have been created through interface redesign of natural oligomers or modular genetic fusions (e.g., Bale et al. 2016, See also reviews and citations therein by Kuhlman and Bradley 2019, Cannon et al. 2020, Vulovic et al. 2021, Lee et al. 2024). More recently, advances in generative deep learning and computational protein design have enabled the *de novo* construction of protein nanoparticles with tailored geometries and tunable symmetries (San Segundo-Acosta et al. 2026). Collectively, these capabilities make protein nanoparticles attractive scaffolds for structural biology (Castells-Graells et al. 2023), vaccine development (Ueda et al. 2020), targeted drug and gene delivery (Hong et al. 2020, Kaltbeitzel and Wich 2023), high-sensitivity medical imaging and theranostics (Kaltbeitzel and Wich 2023), spatial enzyme confinement for biocatalytic nanoreactors (Ren et al. 2019), and size-controlled biomineralization and hierarchical nanomaterials synthesis (Aumiller et al. 2018, Vigil and Spangler 2024). Repeating icosahedral or dodecahedral symmetry supports high-density multivalent display that enhances binding avidity, exemplified by I3-01 and E2p nanoparticles presenting SARS-CoV-2 spike antigens to elicit potent humoral and T-cell responses, while two-component cages such as I53-50 mimic viral capsid geometry (Bale et al. 2016, He et al. 2021). Simultaneously, extensive inter-subunit interfaces render these cages mechanically rigid, making them valuable imaging scaffolds for single-particle cryoEM by adding molecular mass and symmetry that facilitate particle alignment of small attached proteins (Castells-Graells et al. 2023).

Despite these advantages, directly fusing a target protein of interest onto a cage subunit frequently compromises folding and solubility, requiring labor-intensive linker and interface optimization for each new cargo (Antanasijevic et al. 2020). Furthermore, optimal display mechanics differ across applications: whereas cryoEM structural determination requires rigid target immobilization to enable coherent particle averaging, nanoparticle vaccines often benefit from controlled linker flexibility to optimize B-cell receptor cross-linking and antibody breadth (Ellis et al. 2023). These competing demands motivate a modular display architecture in which a standardized, conserved cage core recruits interchangeable targets via engineered binding modules (Liu et al. 2019). Prior scaffolding efforts demonstrated this concept by rigidly fusing designed ankyrin repeat proteins (DARPins) to protein cages (Liu et al. 2019); however, generating target-specific DARPins or antibody fragments traditionally relies on animal immunization or large experimental library selections, creating a bottleneck when pivoting to new targets (Bennett et al. 2023).

Recent breakthroughs in diffusion-based backbone generation and deep-learning sequence design including RFdiffusion, BindCraft, ProteinMPNN, and AlphaFold now enable the rapid *in silico* generation of *de novo* binders against user-defined target epitopes (Jumper et al. 2021, Dauparas et al. 2022, Watson et al. 2023, Pacesa et al. 2025). Because these binders typically terminate in well-ordered α-helices, they are naturally suited for continuous helical fusion to the terminal helices of symmetric cage subunits. Here, we combine *de novo* helical binder design with structure-guided disulfide stapling across the binder–core junction to eliminate lever-arm flexibility, establishing a generalizable platform for high-resolution cryoEM visualization of sub-20 kDa proteins and modular nanoparticle functionalization.

## Results

### End-to-end workflow for generating functionalized protein nanoparticles via fused designed binders

To establish a modular and generalizable display platform, we developed an integrated computational-to-structural pipeline that couples *de novo* helical binder design with rigid helical fusion onto symmetric protein cages (**Fig. 1**). First, given a target protein structure, complementary generative design workflows such as RFdiffusion paired with ProteinMPNN and BindCraft are used to generate candidate α-helical binders against surface hotspots, which are subsequently triaged via AlphaFold confidence filtering and experimental expression/binding assays (**Fig. 1a**). Second, the validated binder is genetically fused to the N-terminal α-helix of a trimeric cage-forming subunit so that the junction forms a continuous, uninterrupted α-helix free of steric clashes with neighboring cage subunits (**Fig. 1b**). Third, the binder–core fusion construct is expressed in *Escherichia coli* (*E. coli*) either as an isolated trimer for *in vitro* assembly with a complementary pentameric building block or via a bicistronic operon for direct *in vivo* self-assembly into 60-subunit icosahedral or dodecahedral nanoparticles (**Fig. 1c**). Finally, incubating the purified, binder-decorated cages with an excess of untagged target protein saturates all 60 surface binding sites, generating monodisperse, high-molecular-weight complexes amenable to cryoEM structure determination and downstream biochemical applications.

**Figure 1.**
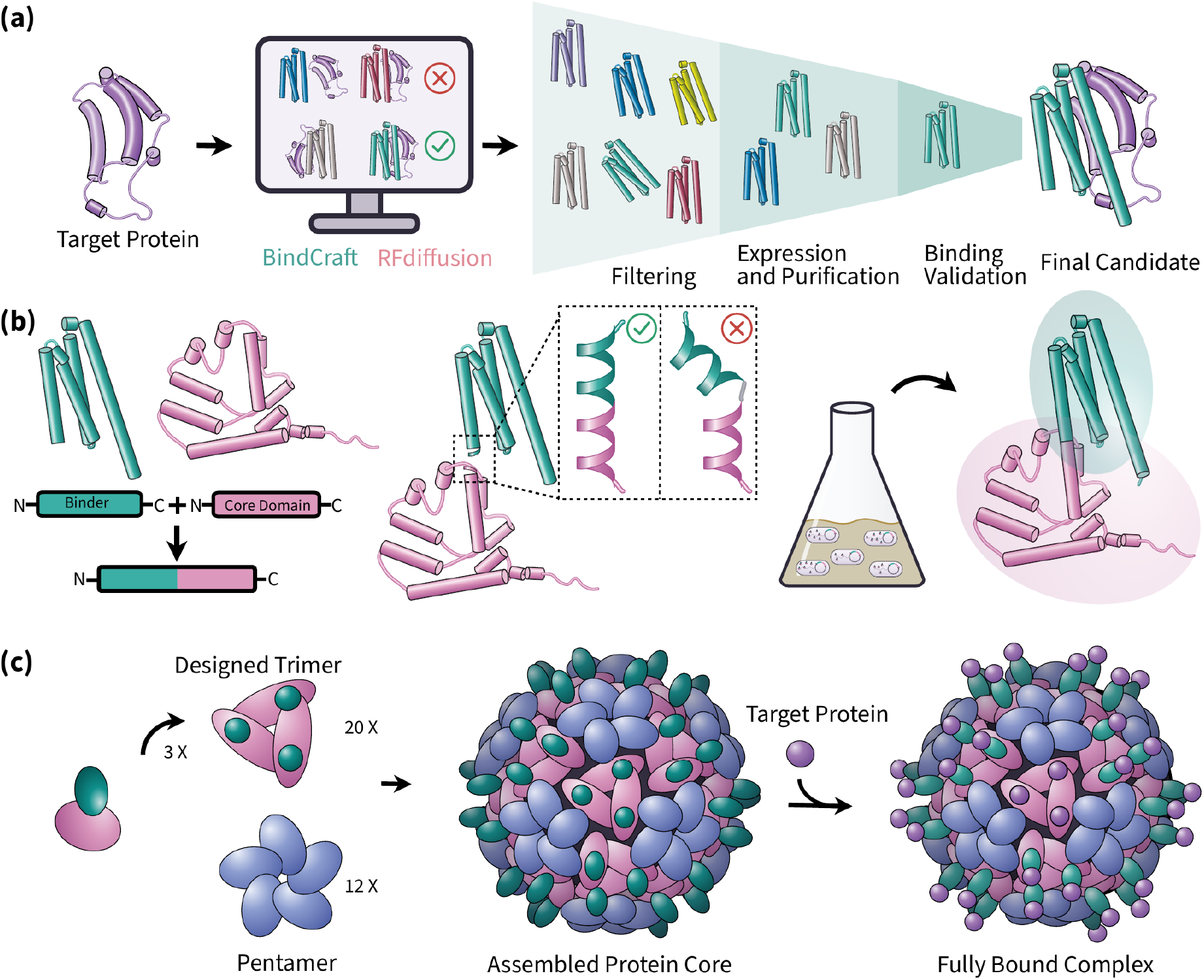
Overview of the modular workflow for displaying target proteins on binder–core assemblies. **(a)** *De novo* computational binder generation using BindCraft or RFdiffusion–ProteinMPNN against the target protein, followed by AlphaFold confidence filtering, recombinant expression/purification, and experimental binding validation. **(b)** Genetic fusion of the validated helical binder to the N-terminus of a trimeric cage subunit via a continuous α-helix engineered to maintain helical stability and prevent steric clashes. **(c)** Self-assembly of trimeric binder–core subunits (alone or with a complementary pentameric building block) dodecahedral or icosahedral nanoparticles, followed by incubation with excess target protein to generate saturated binder–core–target complexes.

### *De novo* design and biochemical characterization of binders against BRD4-BD1

We selected bromodomain 1 of human bromodomain-containing protein 4 (BRD4-BD1) as a stringent proof-of-concept target to evaluate scaffold-assisted cryoEM imaging. BRD4 is an epigenetic transcriptional regulator that drives oncogenic gene expression in acute myeloid leukemia, ovarian carcinoma, and triple-negative breast cancer (Donati et al. 2018, Gilan et al. 2020). Although BRD4-BD1 is a high-priority target for small-molecule and biologics development, its low molecular mass (16 kDa) falls well below the ∼50 kDa practical threshold for conventional single-particle cryoEM, where low signal-to-noise ratios preclude accurate particle orientation alignment (Wentinck et al. 2022).

To generate binders against BRD4-BD1, we executed two parallel computational design campaigns (**Fig. 2a**). In the first approach, RFdiffusion generated 1,000 poly-glycine binder backbones (100–150 residues) targeting the BRD4-BD1 surface, followed by ProteinMPNN-FastRelax sequence design (two sequences per backbone; 2,000 sequences total). Structure prediction of all 2,000 complexes with AlphaFold2 and filtering by strict structural confidence thresholds (complex pLDDT > 0.85, binder pLDDT > 0.70, interface predicted aligned error PAE^interaction^ < 7.0 Å, and binder-aligned RMSD < 3.0 Å) yielded 15 compliant designs, from which the top five candidates were selected. In parallel, BindCraft was run with trajectory-level filtering (complex pLDDT ≥ 0.80, pTM ≥ 0.55, binder pLDDT ≥ 0.80, binder RMSD ≤ 3.5 Å) until five independent passing designs were obtained.

**Figure 2.**
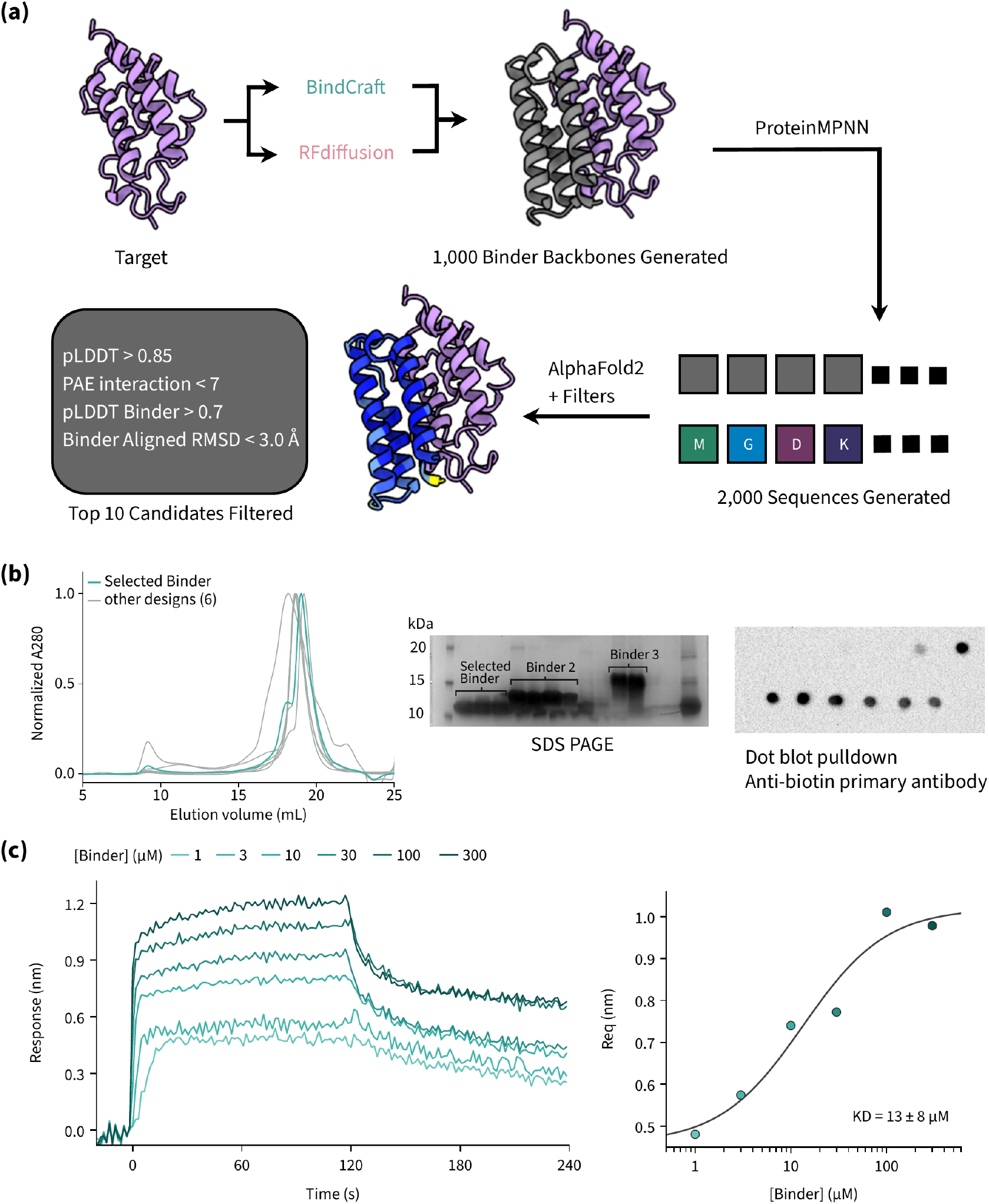
Computational binder generation pipeline and experimental validation against BRD4-BD1. **(a)** Schematic of the *in silico* design workflow against BRD4-BD1: 1,000 backbones were generated with RFdiffusion and assigned two sequences each via ProteinMPNN (2,000 sequences total), followed by AlphaFold2 structure prediction and metric filtering alongside BindCraft generation to select 10 total candidate designs. **(b)** Biochemical characterization of purified binders. *Left:* Overlaid size-exclusion chromatography (SEC) elution profiles of soluble binder designs eluting at 18–20 mL (selected top binder shown in teal). *Middle:* SDS-PAGE analysis of representative purified binder candidates confirming expected molecular weights (∼10-14 kDa). *Bottom right:* Dot-blot pulldown assay against biotinylated BRD4-BD1 probed with an anti-biotin primary antibody, identifying the top-binding candidate. **(c)** Biolayer interferometry (BLI) sensorgram (*left*, 1–300 µM binder) and steady-state equilibrium binding curve (*right*) of the selected binder against immobilized BRD4-BD1, yielding a steady-state K_d_ of 13 ± 8 µM.

All 10 candidate genes were synthesized and expressed in *E. coli* BL21(DE3). Eight of the 10 constructs expressed solubly and eluted as monodisperse monomeric peaks at 18–20 mL on size-exclusion chromatography (SEC), displaying single prominent bands at the expected molecular weight (∼12-14 kDa) on SDS-PAGE (**Fig. 2b**). To screen for target engagement, purified binders were evaluated via dot-blot pulldown assays against biotinylated BRD4-BD1, revealing specific binding for three of the eight soluble designs (**Fig. 2b**). Quantitative kinetic characterization of the top-performing candidate using biolayer interferometry (BLI) across six concentrations (1–300 µM) revealed rapid association and dissociation kinetics with a steady-state dissociation constant (K_d_) of 13 ± 8 µM (**Fig. 2c**). These results confirmed that a compact *in silico* design campaign yields a soluble, helical binder suitable for scaffold integration.

### Biochemical and structural validation of initial continuous-helix binder–core assemblies

To test whether our binder module could generalize across distinct cage geometries, we fused the validated BRD4-BD1 binder via a continuous N-to-C-terminal α-helical junction onto two trimeric building blocks: the single-component dodecahedral pyruvate dehydrogenase complex (PDC) E2p core (PDB: 6CT0) and the trimeric I53-50A subunit of the two-component icosahedral I53-50 nanoparticle (PDB: 7SGE) (**Figs. 1b and 3a**). Both engineered constructs properly oligomerized into trimers and assembled into nanoparticles, eluting at 10–12 mL on a Superose 6 SEC column, consistent with intact cages ∼25 nm in diameter (**Fig. 3b**). Negative-stain transmission electron microscopy (TEM) (**Fig. 3c**) and reference-free cryoEM 2D class averages (**Fig. 3d**) confirmed homogeneous, well-folded spherical assemblies approximately 250 Å in diameter for both scaffolds.

**Figure 3.**
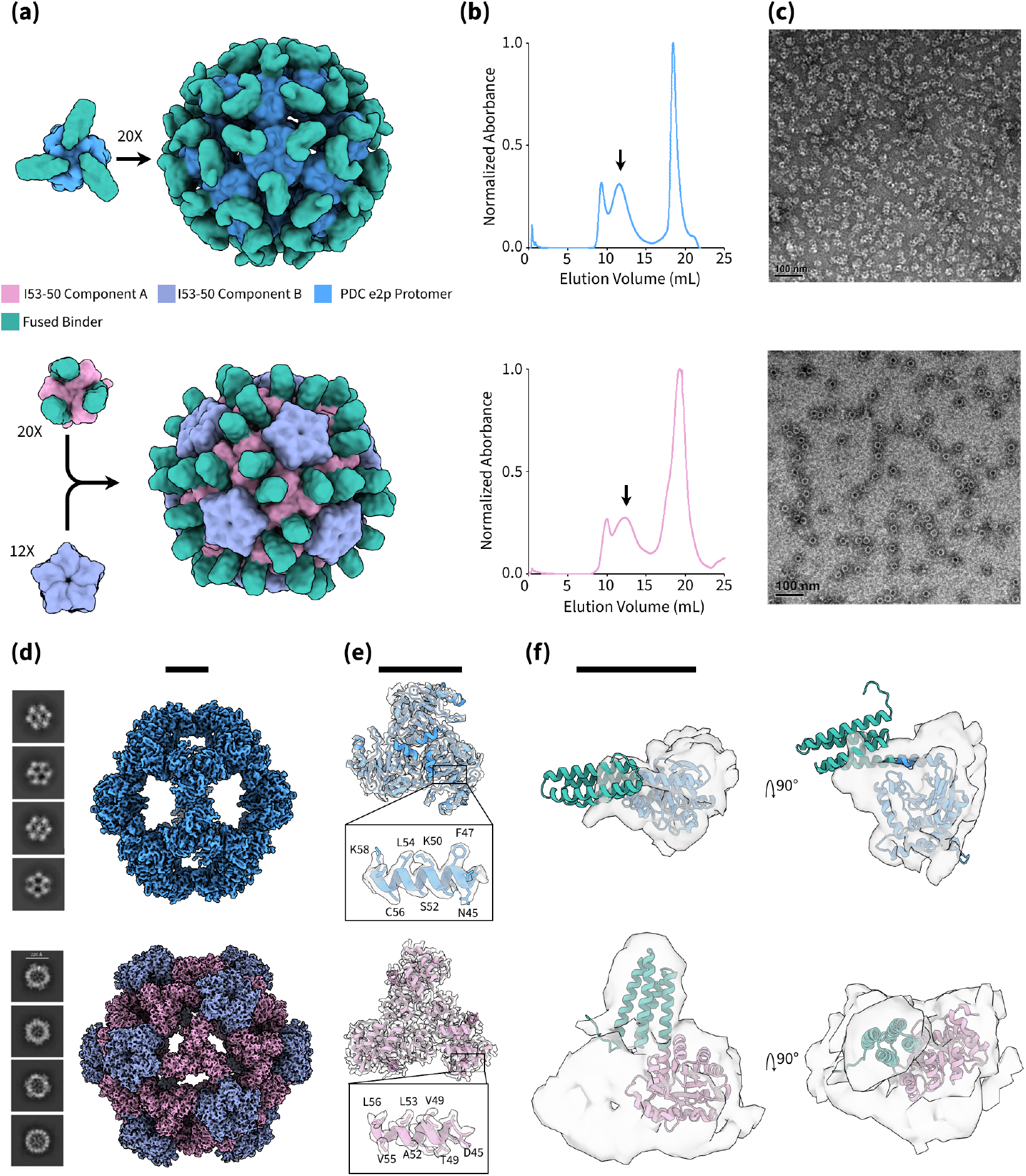
Biochemical and cryoEM validation of initial continuous-helix binder–core assemblies. **(a)** Assembly schematics for binder-fused dodecahedral pyruvate dehydrogenase complex (PDC) E2p (*top*) and two-component icosahedral I53-50 (*bottom*). Core subunits are shown in blue (E2p), pink (I53-50A), and purple (I53-50B); fused binders are shown in teal. **(b)** Size-exclusion chromatography (SEC) profiles of assembled E2p (*top*) and I53-50 (*bottom*) nanoparticles eluting at 10–12 mL (arrows). **(c)** Representative negative-stain transmission electron micrographs of purified E2p (*top*) and I53-50 (*bottom*) nanoparticles. Scale bars, 100 nm. **(d)** Reference-free cryoEM 2D class averages and icosahedral 3D reconstructions of the E2p (2.97 Å, *top*) and I53-50 (2.34 Å, *bottom*) cores. Scale bars, 50 Å. **(e)** CryoEM density maps of the trimeric building blocks with enlarged views of representative core α-helices showing well-resolved side-chain densities. Scale bars, 50 Å. **(f)** Representative focused cryoEM 3D classification maps showing low-resolution, poorly defined peripheral density for the fused binders in both E2p (*top*) and I53-50 (*bottom*), illustrating lever-arm flexibility at the unstapled helical junction. Scale bars, 50 Å.

We next determined whether the continuous α-helical linker alone provided sufficient conformational rigidity for high-resolution cryoEM reconstruction of the displayed binder. Single-particle cryoEM reconstructions with icosahedral symmetry yielded high-resolution maps of the underlying cores at 2.97 Å for PDC E2p and 2.34 Å for I53-50, clearly resolving backbone carbonyls and bulky side chains within the core helices (**Figs. 3d and 3e**). In contrast, density corresponding to the fused binder was severely attenuated in both reconstructions and only appeared at low resolution (∼15 Å) in focused 3D classifications (**Fig. 3f**). For PDC E2p, only the proximal segment of the fusion helix extending from the core was discernible; for I53-50, diffuse density for the four-helix bundle was visible, but focused classification and local refinement failed to improve map interpretability. This behavior is indicative of a geometric lever-arm effect: because both the cage subunit and the binder are globular domains connected solely by an unsupported α-helix, sub-nanometer angular bending or torsional flexing at the junction translates into multi-angstrom positional displacement at the distal tip of the binder. Moreover, because PDC E2p fusions exhibited lower expression yields, incomplete maltose-binding protein (MBP) tag cleavage, and reduced solubility during scale-up, we selected the high-yielding bicistronic I53-50 platform for subsequent interface rigidification.

### Disulfide stapling of the binder–core junction suppresses flexibility and enables near-atomic binder visualization

To eliminate angular play at the fusion junction without altering the target-binding interface, we inspected the AlphaFold3-predicted trimeric structure of the I53-50A–binder fusion and identified two spatially proximal residues, C104 on the binder fusion helix and C213 on an adjacent helix of the I53-50A core domain, whose Cβ–Cβ geometry was compatible with an engineered disulfide bond (**Fig. 4a**). Following identification of the C104/C213 cysteine pair, we applied ProteinMPNN to redesign neighboring residues along the fusion helix (positions 99–112) to pack favorably around the disulfide staple and preserve helical propensity.

**Figure 4.**
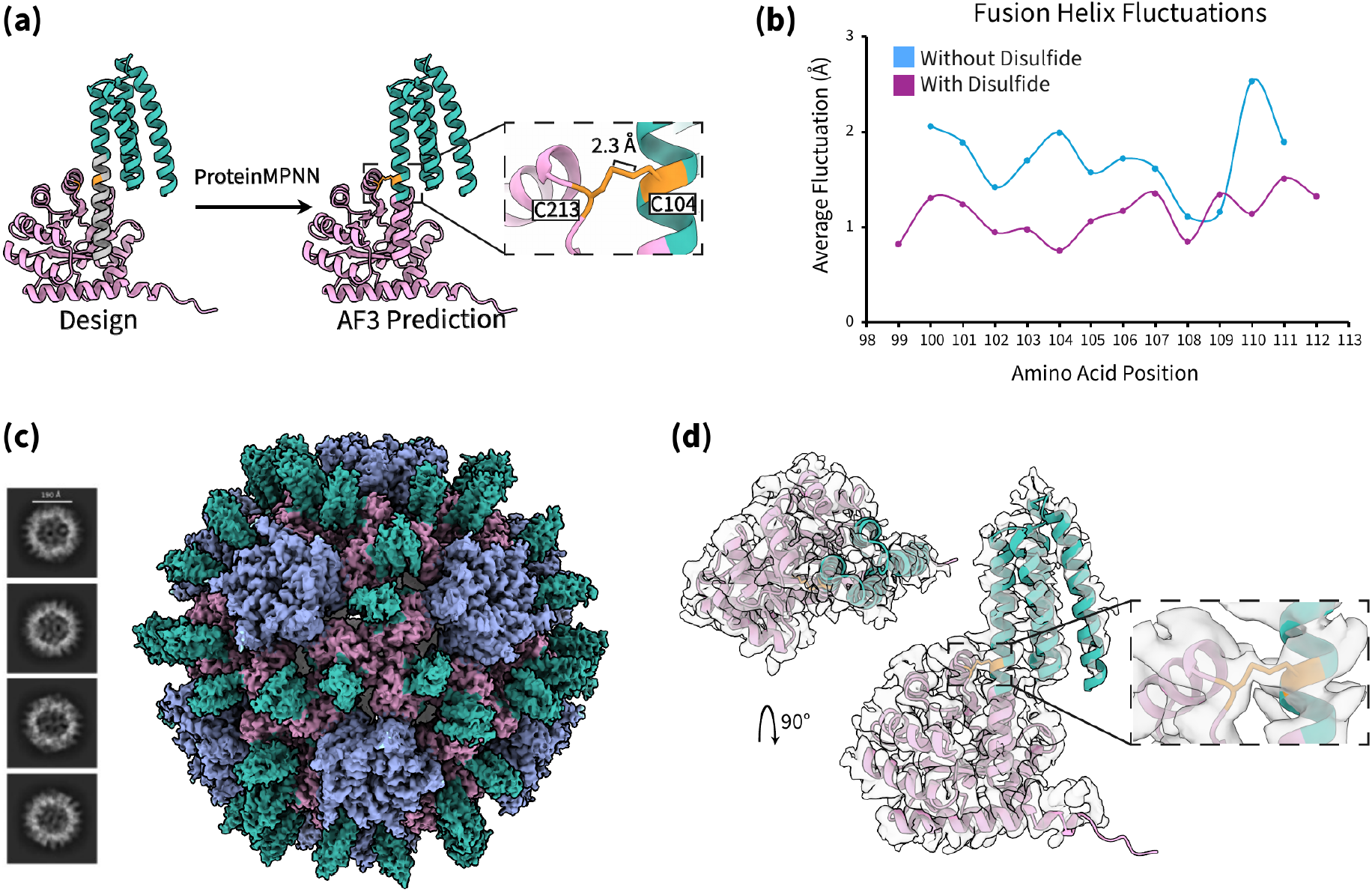
Disulfide stapling of the I53-50 binder–core junction suppresses flexibility and enables high-resolution binder reconstruction. **(a)** Structure-guided engineering of an inter-domain disulfide bond (C104–C213) bridging the binder fusion helix (teal) and the I53-50A core domain (pink), followed by ProteinMPNN sequence redesign of surrounding junction residues and AlphaFold3 validation (predicted S–S distance, 2.3 Å). **(b)** All-atom molecular dynamics (100 ns) backbone root-mean-square fluctuation (RMSF) profiles across the fusion helix (residues 98–113) before (blue) and after (purple) introduction of the C104–C213 disulfide bond. **(c)** Reference-free cryoEM 2D class averages (*left*, scale bar 150 Å) and icosahedral 3D reconstruction (*right*) of disulfide-stapled I53-50–binder nanoparticles, revealing well-ordered radial binder protrusions at all 60 sites. **(d)** Locally refined 3.2 Å cryoEM density map (semi-transparent surface) of a single binder–I53-50A protomer fitted with the AlphaFold3 atomic model (I53-50A in pink, binder in teal, engineered C104–C213 disulfide bond in orange/yellow). *Inset:* Close-up of the fusion junction showing continuous cryoEM density across the engineered disulfide bridge.

Before experimental production, we evaluated the dynamic impact of the disulfide staple using all-atom molecular dynamics (MD) simulations in GROMACS (**Fig. 4b**). Across 100 ns production trajectories in explicit solvent, the disulfide-stapled construct exhibited markedly reduced backbone mobility across the fusion junction (residues 98–113), lowering per-residue root-mean-square fluctuations (RMSF) by 0.2 to 1.4 Å relative to the unstapled continuous-helix construct (where fluctuations peaked above 2.5 Å at residue 110).

Encouraged by these *in silico* stabilization metrics, we expressed and purified the disulfide-stapled I53-50–binder nanoparticles via bicistronic co-expression in *E. coli*. Strikingly, cryoEM 2D class averages and icosahedral 3D reconstructions revealed well-defined, spike-like protrusions radiating symmetrically from all 60 trimeric positions on the I53-50 surface. This was a stark contrast to the blurred, featureless exterior of the unstapled construct (**Fig. 4c** vs. **Fig. 3f**). Symmetry expansion followed by focused 3D classification and local refinement of a single asymmetric binder–core protomer yielded a 3.2 Å density map that resolved the entire four-helix bundle of the *de novo* binder (**Fig. 4d**). Rigid-body docking of the AlphaFold3 model confirmed continuous density bridging C104 and C213 at an S–S bond distance (∼2.3 Å in the starting model) consistent with quantitative disulfide bond formation (**Fig. 4d**, inset).

### CryoEM structure of the disulfide-stapled binder–core assembly bound to BRD4-BD1

With the fused binder rigidly anchored on the I53-50 scaffold, we incubated the disulfide-stapled nanoparticles with a molar excess of purified BRD4-BD1 to saturate the surface binding sites prior to vitrification. Even prior to focused classification, an initial icosahedral reconstruction displayed robust, bi-lobed peripheral density at all 60 sites corresponding to both the four-helix binder and bound BRD4-BD1, demonstrating high-occupancy target capture across the nanoparticle surface (**Fig. 5a**).

**Figure 5.**
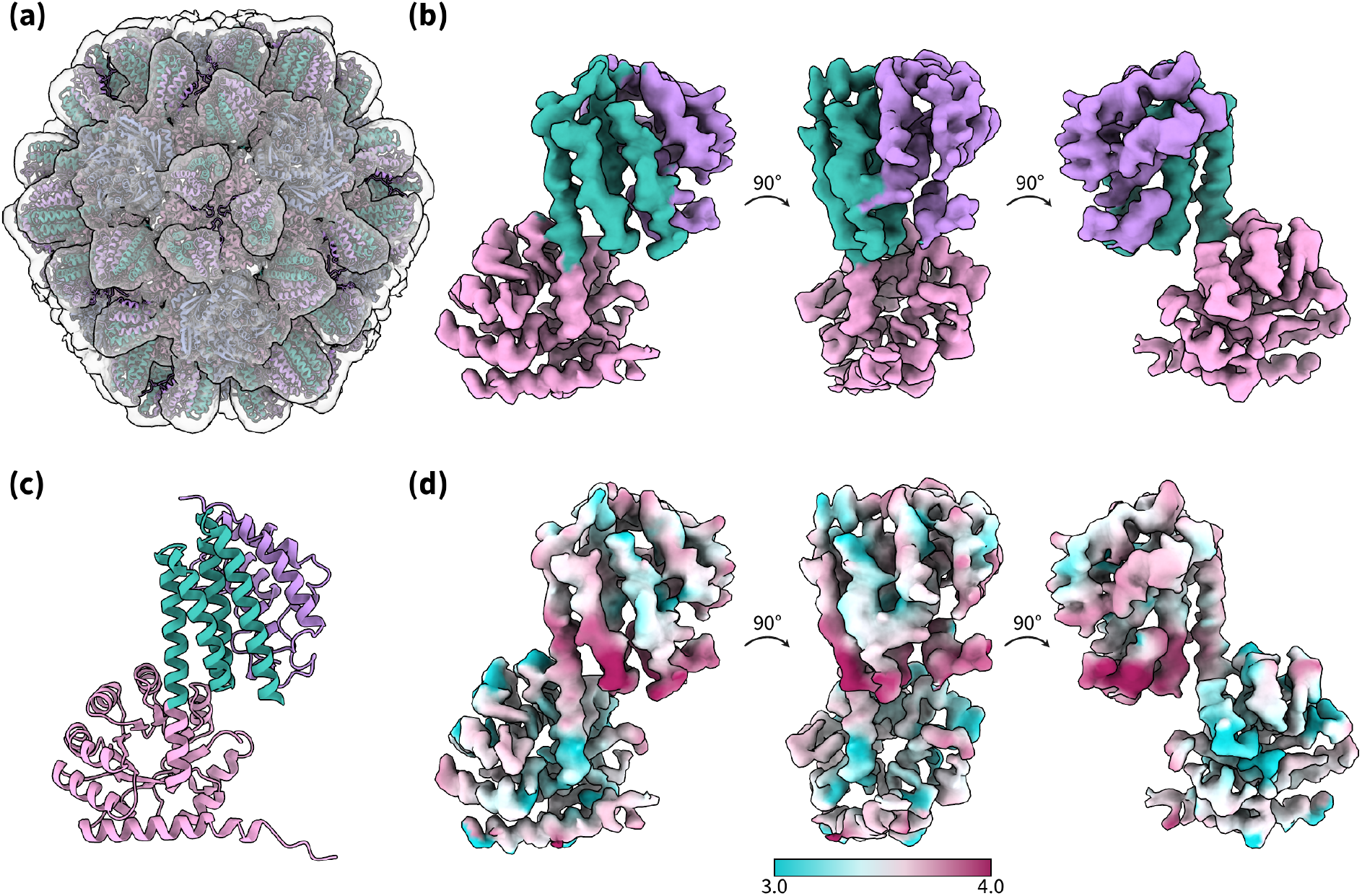
CryoEM structure of the disulfide-stapled I53-50–binder nanoparticle bound to BRD4-BD1. **(a)** Icosahedral homogeneous refinement cryoEM map (transparent envelope fitted with atomic coordinates) showing full occupancy of BRD4-BD1 (purple) bound to the displayed binders (teal) across all 60 binding sites of the I53-50 cage. **(b)** Orthogonal views of the locally refined 3.70 Å cryoEM density map of a single asymmetric binder–core–BRD4-BD1 complex colored by subunit (I53-50A in pink, binder in teal, BRD4-BD1 in purple). **(c)** AlphaFold3-predicted ternary complex model rigid-body docked into the 3.70 Å cryoEM density map (map-to-model correlation = 0.69). **(d)** Local resolution distribution (3.0–4.0 Å) mapped onto the locally refined cryoEM reconstruction shown in **(b)**.

To overcome subtle residual asymmetric motion among the 60 protomers, we performed icosahedral symmetry expansion (generating 1,304,040 asymmetric unit particles from 21,734 intact cages), followed by masked 3D classification, local refinement, and 3D flexible refinement on a single binder–core–BRD4-BD1 protomer. Focused 3D classification isolated a homogeneous class comprising 37% of the symmetry-expanded particles with full BRD4-BD1 occupancy, which refined to an overall resolution of 3.70 Å (**Fig. 5b**), with local resolutions spanning 3.0 Å across the I53-50A core and fusion junction to ∼4.0 Å at the distal solvent-exposed surface of BRD4-BD1 (**Fig. 5d**).

Rigid-body docking of the AlphaFold3-predicted ternary complex into the 3.70 Å cryoEM map using ChimeraX yielded a map-to-model correlation coefficient of 0.69 (**Fig. 5c**). Aside from flexible terminal loops and the N-terminal purification tag, all canonical α-helices of the binder and the four-helix bromodomain fold of BRD4-BD1 were unambiguously accounted for by the experimental density. Importantly, BRD4-BD1 engages the exact surface patch of the binder predicted by the initial RFdiffusion/AlphaFold design model, validating the atomic accuracy of the computational design pipeline. Resolving a 16 kDa protein at 3.70 Å, comparable to the smallest cargoes visualized on any cryoEM imaging scaffold to date (Yi et al. 2026), demonstrates that disulfide-stapled *de novo* helical binders provide an effective route to structural characterization of small, monomeric therapeutic proteins.

## Discussion

In this study, we demonstrate that coupling a *de novo* computationally designed α-helical binder to an icosahedral protein cage via a disulfide-stapled helical junction overcomes the conformational flexibility that has historically limited modular cryoEM imaging scaffolds. Whereas a continuous α-helical fusion supported efficient assembly of both PDC E2p and I53-50 nanoparticles, the displayed binder remained poorly resolved beyond the first turn of the junction helix. Introducing a single engineered disulfide bond (C104–C213) across the binder–core interface, paired with local sequence optimization via ProteinMPNN, suppressed junction fluctuations and enabled near-atomic cryoEM reconstructions of both the apo binder (3.2 Å) and the 16 kDa BRD4-BD1-bound complex (3.70 Å). Because the sole structural prerequisite is a terminal α-helix on the binder, a ubiquitous feature of *de novo* binders and many natural protein domains, this disulfide-stapling blueprint offers a broadly transferable strategy for rigid cargo presentation.

Conformational heterogeneity due to residual flexibility at helical junctions is a well-recognized bottleneck in protein scaffold engineering. Pioneering DARPin-cage scaffolds utilized continuous α-helical fusions to constrain cargo orientation, yet residual flexing along the linker restricted achievable resolution (Liu et al. 2018). Geometrically, any unsupported helix acts as a lever arm in which sub-degree angular deviations at the base amplify linearly with radial distance from the cage center (Camille et al. 2025). Prior efforts have addressed this challenge by engineering secondary inter-trimer contacts between adjacent DARPins (Castells-Graells et al. 2023), embedding targets within dimeric coiled-coil stalks (Camille et al. 2025), or introducing inter-chain disulfides into Fab elbow regions (Kung et al. 2025). Our strategy unites the simplicity of terminal helical fusion with covalent junction stapling: because the disulfide staple bridges the conserved cage subunit and the proximal fusion helix rather than the target-binding interface, the cage-junction module remains invariant when swapping in new binders designed for different targets.

Integrating generative computational binder design into this scaffold architecture also eliminates a major experimental bottleneck. Traditional affinity proteins such as monoclonal antibodies, Fabs, nanobodies, and library-selected DARPins require months of immunization or high-throughput display screening (Bennett et al. 2023). By contrast, diffusion-and deep-learning-based pipelines generate bespoke helical binders against defined epitopes within hours. While current workflows perform best when an experimental structure of the target is available, emerging generative models capable of targeting AlphaFold-predicted structures or conformationally dynamic regions will further expand the range of accessible targets.

Two practical considerations from our proof-of-concept study highlight clear avenues for optimization. First, our modest binder affinity (13 µM) necessitated incubating the scaffold with excess BRD4-BD1 (∼300 µM) to achieve full site saturation, and 3D classification indicated ∼37% occupancy in the highest-resolution particle subset. Because we screened only 10 designs (five each from RFdiffusion and BindCraft), low-to-mid micromolar affinity reflects limited sampling depth rather than a fundamental limit of the scaffold. Scaling experimental screening to 50–100 designs or leveraging next-generation binder design architectures such as BoltzGen and RFD4-Proteina, which routinely achieve low-nanomolar to picomolar affinities (Stark et al. 2025, Avery et al. 2026, Corley et al. 2026), will enable stoichiometric target capture at low concentrations and near-100% particle occupancy. Second, although disulfide stapling improved resolution to 3.70 Å, a modest radial resolution gradient (3.0 Å at the junction to 4.0 Å at the distal target tip) persisted due to the extended distance between the binder and the I53-50A core, which is constrained by steric clearance around the neighboring I53-50B pentamers. Transplanting this disulfide-stapled junction onto single-component porous cages such as I3-01, which shares the trimeric aldolase fold of I53-50A but lacks pentameric caps (Bale et al. 2016), would allow the binder to be recessed closer to the cage surface, further minimizing lever-arm motion and accommodating larger multi-domain cargoes.

Beyond single-particle cryoEM illustrated in this study, the modular and reversible nature of our disulfide-stapled binder–cage platform unlocks broad applications across many fields, mostly notably structural cell biology, therapeutics, and vaccine engineering. First, in cellular cryogenic electron tomography (cryoET), identifying specific macromolecular complexes within the crowded eukaryotic cytoplasm remains a major hurdle. Because our binder–cage assemblies are genetically encodable, self-assembling, and exhibit an unmistakable ∼25 nm symmetric signature with high image contrast, they could be expressed intracellularly or delivered via transfection as genetically encoded molecular fiducials to localize and orient endogenous, untagged target proteins *in situ* (Fung et al. 2023, Luo et al. 2026). Second, unlike permanent backbone fusions, an engineered disulfide staple functions as a reversible redox-responsive switch: disulfides remain intact in the oxidizing extracellular space, preserving rigid multivalent presentation during circulation or lymph-node trafficking, but are selectively reduced in the reducing cytosol, a mechanism widely exploited in antibody–drug conjugates (Pillow et al. 2016). This redox sensitivity could enable triggered intracellular cargo release or allow controlled *in vitro* reduction (e.g., via mild TCEP or DTT treatment) to toggle nanoparticles between a rigid state, for structural determination, and a flexible state, to tune B-cell receptor engagement and antibody breadth in multivalent vaccines (Ellis et al. 2023). Finally, high-avidity 60-copy binder cages immobilized on size-exclusion or membrane filters could serve as high-capacity affinity capture reagents to enrich low-abundance, untagged native proteins or endogenous complexes directly from cell lysates under gentle elution conditions. PDC E2p remains an attractive scaffold for applications where rigidity is not the objective. Its inter-trimer contacts are native rather than designed and lack a second component as in I53-50, giving its trimer-trimer contacts greater conformational freedom. The core is also derived from an essential and highly conserved human enzyme, which is advantageous where biocompatibility matters.

## Materials and Methods

### BRD4-BD1 binder design and expression

The crystal structure of human BRD4-BD1 was preprocessed using PDBFixer to restore missing side-chain atoms and resolve chain discontinuities. *De novo* binders were generated using two complementary workflows. First, RFdiffusion (v1.1.0-42) (Watson et al. 2023) was used to generate 1,000 poly-glycine binder backbones (100–150 amino acids) targeting the BRD4-BD1 surface using default noise scale parameters (noise_scale_ca = 1, noise_scale_frame = 1). Two sequences per backbone (2,000 sequences total) were designed using ProteinMPNN-FastRelax (Dauparas et al. 2022) and predicted in complex with BRD4-BD1 using AlphaFold2 (Jumper et al. 2021). Designs were filtered using a custom Python script requiring complex pLDDT > 0.85, binder pLDDT > 0.70, interface PAE interaction < 7.0 Å, and binder-aligned RMSD < 3.0 Å; 15 sequences passed all filters, and the top five ranked by interface PAE were selected. Second, BindCraft (Pacesa et al. 2025) was executed with standard trajectory filters (complex pLDDT ≥ 0.80, pTM ≥ 0.55, binder pLDDT ≥ 0.80, binder RMSD ≤ 3.5 Å) until five compliant designs were obtained.

Codon-optimized genes encoding all 10 binders with an N-terminal 6×His tag in pET-29b(+) were synthesized (Twist Bioscience) and transformed into *E. coli* BL21(DE3) (Thermo Fisher Scientific). Starter cultures (25 mL LB with 50 µg/mL kanamycin) were grown overnight at 37 °C (225 rpm) and used to inoculate 1 L Terrific Broth (TB) supplemented with 50 µg/mL kanamycin. Cultures were grown at 37 °C to an OD^600^ of 1.0 and induced with 1 mM IPTG at 18 °C for 18 h. Harvested cell pellets were resuspended in lysis buffer (50 mM Tris-HCl pH 7.5, 300 mM NaCl, 5% [v/v] glycerol) supplemented with Benzonase, lysozyme, and EDTA-free protease inhibitor tablets (Thermo Fisher Scientific), and lysed via high-pressure homogenization (Emulsiflex). Clarified lysates were incubated with immobilized cobalt affinity resin (Thermo Fisher Scientific) for 30 min at room temperature, washed, and eluted with lysis buffer containing 500 mM imidazole. Eluates were concentrated to 500 µL using 3 kDa MWCO Amicon Ultra centrifugal filters (Millipore) at 4,000 × g and purified by size-exclusion chromatography (SEC) on an ÄKTA pure system (Cytiva).

### Biolayer interferometry (BLI)

BLI binding assays were performed on an Octet QK384 instrument (Sartorius, formerly ForteBio) at 30 °C with orbital shaking at 1,000 rpm using streptavidin (SA) biosensors (Sartorius) in kinetics buffer (1× PBS pH 7.4, 0.1% [w/v] BSA, 0.02% [v/v] Tween-20). Following a 60 s initial baseline, biotinylated BRD4-BD1 (10 µg/mL) was immobilized onto SA sensors to a defined response threshold (∼0.5–1.0 nm) rather than for a fixed duration, ensuring comparable ligand density across sensors. Loaded sensors were washed and equilibrated in a second baseline (≥120 s) in kinetics buffer, then dipped into a serial dilution series of purified binder (1, 3, 10, 30, 100, and 300 µM) for association (120 s) followed by dissociation (120 s) in kinetics buffer. Binder dilutions were prepared in glycerol-free kinetics buffer to avoid bulk refractive-index artifacts, and each sensor was used for a single association–dissociation cycle without regeneration.

Double-reference subtraction was performed using a ligand-loaded sensor exposed to buffer alone (baseline/drift reference) and an unloaded sensor exposed to the highest binder concentration (300 µM; non-specific/bulk reference). Because association and dissociation were too rapid to resolve reliable rate constants, affinity was determined by steady-state analysis: the equilibrium response (Req) at each concentration was plotted against binder concentration and fit to a 1:1 binding isotherm. A concentration-independent response offset term was included in the steady-state fit to account for a non-specific signal observed at high binder concentrations; omission of this term yielded a substantially different apparent affinity and a poor fit to the data. Fits were performed in Sartorius (ForteBio) Data Analysis software (version 11.0.0.50) and independently verified using a custom Python script developed for this analysis. The reported Kd values are determined from the optimized assay configuration (threshold-controlled ligand loading, double-reference subtraction, and offset-corrected steady-state fitting); an initial pilot experiment performed under higher ligand-loading conditions yielded a consistent order-of-magnitude estimate, supporting the robustness of the measurement.

### AlphaFold structure predictions and molecular dynamics simulations

Monomeric and trimeric structure predictions were generated using the AlphaFold Server running AlphaFold3 (Jumper et al. 2021), and top-ranked models were used for disulfide site identification, MD simulations, and cryoEM map docking. For MD simulations, trimeric AlphaFold3 models of the binder fused to I53-50A (with and without the C104–C213 disulfide bond) were solvated in rectangular TIP3P water boxes (10 Å padding) neutralized with 150 mM KCl using CHARMM-GUI Solution Builder (Jo et al. 2008) with the CHARMM36m force field. Simulations were executed in GROMACS (Abraham et al. 2015) under periodic boundary conditions: systems were energy-minimized, equilibrated at 298 K for 100 ps, and simulated for 100 ns of unrestrained production dynamics. Backbone RMSD and per-residue RMSF across the fusion helix (residues 98–113) were computed to quantify junction rigidity.

### BRD4-BD1 expression, *in vivo* biotinylation, and purification

A pET-29b(+) plasmid encoding human BRD4-BD1 with an N-terminal 6×His tag, HRV 3C/TEV cleavage sites, and an AviTag (GLNDIFEAQKIEWHE) was co-transformed with a pET-22b(+) vector encoding BirA biotin ligase into *E. coli* BL21(DE3). Cultures were grown in TB supplemented with 50 µg/mL kanamycin and 50 µg/mL ampicillin, and supplemented with 100 µM D-biotin upon induction with 1 mM IPTG at 18 °C. Biotinylated BRD4-BD1 was purified by cobalt affinity chromatography and SEC as described above.

### Cloning and assembly of I53-50 and PDC E2p binder–core nanoparticles

For I53-50 constructs, the selected binder was fused via a continuous α-helix to the N-terminus of the trimeric I53-50A subunit (PDB: 7SGE, chain A) with an N-terminal 6×His tag. Gene fragments (Twist Bioscience) and pET-29b(+) vector backbones were PCR-amplified and assembled using NEBuilder HiFi DNA Assembly Master Mix (New England Biolabs). Pentameric I53-50B was cloned separately into pET-29b(+). For *in vitro* assembly, I53-50A–binder and I53-50B were expressed separately, purified, and mixed at a 1:1 molar ratio (Bale et al. 2016). To streamline production via *in vivo* assembly, a bicistronic pET-29b(+) plasmid co-expressing both I53-50A–binder and I53-50B was constructed via Gibson Assembly and transformed into *E. coli* BL21(DE3) pLysS (Thermo Fisher Scientific). Cultures were grown with 50 µg/mL kanamycin and 34 µg/mL chloramphenicol, induced at 18 °C, and lysed in CHAPS-supplemented buffer (50 mM Tris-HCl pH 7.5, 300 mM NaCl, 0.75% [w/v] CHAPS, 5% [v/v] glycerol) to prevent cage precipitation. Assembled nanoparticles were purified by cobalt affinity chromatography, concentrated in 100 kDa MWCO Amicon Ultra filters, and isolated on a Superose 6 Increase 10/300 GL column (Cytiva) at 0.5 mL/min.

For dodecahedral PDC E2p constructs, the binder was fused to the N-terminus of the E2p core protomer (PDB: 6CT0) downstream of an N-terminal 6×His–MBP solubility tag and TEV protease cleavage site. Following cobalt affinity purification, overnight incubation with TEV protease (1:1 mass ratio) yielded incomplete MBP cleavage and low soluble nanoparticle yields, prompting focus on I53-50.

### SDS-PAGE, dot-blot pulldown, and negative-stain electron microscopy

For SDS-PAGE, 3 µL fractions were mixed with 10 µL lysis buffer, 4 µL LDS sample buffer, and 0.88% (v/v) β-mercaptoethanol, heated at 95 °C for 3 min, and resolved on 4–20% Mini-PROTEAN TGX gels (Bio-Rad) at 130 V for 50 min alongside an 11–270 kDa protein ladder (HelloBio), followed by InstantBlue staining (Abcam). For dot-blot binding assays, 3 µL protein samples were spotted onto methanol-activated nitrocellulose membranes, air-dried, blocked with 5% (w/v) non-fat milk in TBST (TBS, 0.1% Tween-20) for 1 h, incubated with mouse anti-biotin primary antibody (Thermo Fisher Scientific, 1 h) and HRP-conjugated anti-mouse secondary antibody (1 h), and developed with Immobilon HRP substrate (Millipore Sigma) on a ChemiDoc imager (Bio-Rad).

For negative-stain TEM, SEC-purified binder–cores (200 µg/mL, 3 µL) were applied to glow-discharged carbon/formvar copper grids (PELCO) for 1 min, blotted, stained three times with 3 µL of 2% (w/v) uranyl acetate (Electron Microscopy Sciences) with a 30 s final incubation, air-dried, and imaged on a Tecnai T12 microscope at 26,000× magnification.

### CryoEM sample preparation, data acquisition, and image processing

Purified disulfide-stapled I53-50–binder nanoparticles (1 mg/mL) were incubated with BRD4-BD1 (5 mg/mL) for 30 min at room temperature. Aliquots (3 µL) were applied to glow-discharged Quantifoil R1.2/1.3 copper grids overlaid with a continuous graphene oxide film, blotted for 10 s at 4 °C and 100% relative humidity (blot force 5), and plunge-frozen into liquid ethane/propane using a Vitrobot Mark IV (Thermo Fisher Scientific). Movies (10,303 total) were recorded on a Titan Krios G4 transmission electron microscope (Thermo Fisher Scientific) operated at 300 kV equipped with a Falcon 4i direct electron detector at 165,000× nominal magnification (0.756 Å/pixel) in EER format with a total electron dose of 43 e^−^/Å^2^.

Image processing was performed in CryoSPARC v4.7.1 (Punjani et al. 2017). After patch motion correction and patch CTF estimation, particles were picked via blob and template pickers and extracted with 8× binning. Following iterative 2D classification, 21,734 intact cage particles were selected for *ab initio* reconstruction and homogeneous refinement with icosahedral (I) symmetry. Icosahedral symmetry expansion generated 1,304,040 asymmetric protomer sub-particles. A soft mask (threshold 0.005, dilation radius 4 pixels, soft padding 12 pixels) encompassing one I53-50A–binder–BRD4-BD1 protomer was applied for focused 3D classification without alignment. A single well-resolved class containing 37% of the symmetry-expanded sub-particles exhibited full BRD4-BD1 density and was subjected to local refinement and 3D flexible refinement to yield a final 3.70 Å reconstruction. Atomic models predicted by AlphaFold3 were rigid-body docked into the cryoEM maps using UCSF ChimeraX with minor adjustments needed manually done in Coot (Emsley et al. 2010).

## Author Contributions

**Roman Aguirre**: Writing – original draft; writing – review and editing; visualization; investigation; data curation, formal analysis; supervision. **Hongbo Zhu:** Writing – original draft; writing – review and editing; visualization; investigation; data curation. **Todd Schurr**: Investigation; data curation; formal analysis, writing; editing. **Robert Damoiseaux**: Investigation; formal analysis, writing; editing. **Z. Hong Zhou**: Funding acquisition; Supervision; project administration; conceptualization; writing – review and editing

## Acknowledgements

This work was supported in part by grants from the National Institutes of Health (R01GM071940 to ZHZ and its diversity supplement to RA). We acknowledge the Electron Imaging Center for Nanosystems (EICN; RRID:SCR_022900) and the Molecular Screening Shared Resource (MSSR; RRID:SCR_022956) at the California NanoSystems Institute (CNSI), UCLA, for instrumentation, screening support, and technical assistance. CryoEM data were acquired using the Titan Krios G1 (supported by NIH S10RR23057) and Titan Krios G4 (supported by NIH S10OD032459). We thank Mark Arbing for guidance, instrument access, and providing biotinylated BRD4-BD1.

