## Supplementary Material for "Generalizable strategy for designing helical binders rigidly fused to large protein cages with disulfide bonds"

### Supplemental Data

#### Binder “F1” sequence:

MHHHHHHMEELVESMKYAAEKNLKNVKELAEVAKKKEQAKTALKAYIKGSVSTMIEAVKATKNKEMAEIVKKM
VEEIKKIAEDISEEVKKVAEEMYEKVEKVMKEIE*

#### I53-50A sequence with binder (disulfide bond engineered):

MHHHHHHEELVESMKYAAEKNLKNVKELAEVAKKKEQAKTALKAYIKGSVSTMIEAVKATKNKEMAEIVKKMVE
EIKKIAEDISEEVKKVAEEMYEKVEKEI**C**NLEAKEEFEKALEKEKIVAVLRANSVEEAIEKAVAVFAGGVHLIEITFT
VPDADTVIKALSVLKEKGAIIGAGTVTSVEQCRKAVESGAEFIVSPHLDEEISQFCKEK**C**VFYMPGVMTPTELVK
AMKLGHDILKLFPGEVVGPEFVKAMKGPFPNVKFVPTGGVDLDNVCEWFDAGVLAVGVGDALVEGDPDEVRE
KAKEFVEKIRGCTEGSLEWSHPQFEK*

*Highlighted cysteine residues (C104 and C213) denote the engineered disulfide.*

#### I53-50A sequence with binder (no disulfide bond):

MHHHHHHEELVESMKYAAEKNLKNVKELAEVAKKKEQAKTALKAYIKGSVSTMIEAVKATKNKEMAEIVKKMVE
EIKKIAEDISEEVKKVAEEMYEKVEKVMKEEEAARKMEELFKKHKIVAVLRANSVEEAIEKAVAVFAGGVHLIEITF
TVPDADTVIKALSVLKEKGAIIGAGTVTSVEQCRKAVESGAEFIVSPHLDEEISQFCKEKGVFYMPGVMTPTELV
KAMKLGHDILKLFPGEVVGPEFVKAMKGPFPNVKFVPTGGVDLDNVCEWFDAGVLAVGVGDALVEGDPDEVR
EKAKEFVEKIRGCTEGSLEWSHPQFEK*

#### I53-50B sequence:

MNQHSHKDHETVRIAVVRARWHAEIVDACVSAFEAAMRDIGGDRFAVDVFDVPGAYEIPLHARTLAETGRYGA
VLGTAFVVNGGIYRHEFVASAVINGMMNVQLNTGVPVLSAVLTPHNYDKSKAHTLLFLALFAVKGMEAARACVE
ILAAREKIAAGSLE*

#### BRD4-BD1 sequence:

MHHHHHHLEVLFQGPGLNDIFEAQKIEWHEENLYFQSGGSNPPPPETSNPNKPKRQTNQLQYLLRVVLKTLW
KHQFAWPFQQPVDAVKLNLPDYYKIIKTPMDMGTIKKRLENNYYWNAQECIQDFNTMFTNCYIYNKPGDDIVLM
AEALEKLFLQKINELPTEET*
